# Enhanced processing of cartoons in infant visual cortex

**DOI:** 10.64898/2026.05.11.724345

**Authors:** Tristan S. Yates, Ariadne Letrou, Juliana E. Trach, Lillian Behm, Dawoon Choi, Cameron T. Ellis, Nicholas B. Turk-Browne

## Abstract

Developing sensory systems may have heightened sensitivity to exaggerated features that emphasize diagnostic information, as shown by the benefits of infant-directed speech for language acquisition. Here, we examine this sensory exaggeration hypothesis in the visual domain by testing whether cartoons elicit stronger and more consistent neural representations than realistic movies in human infants. We collected fMRI data while 24 awake infants (4–15 months) watched the same 3-minute clip of the opening sequence from the original animated version of The Lion King (1994) and its shot-for-shot remake with photorealistic CGI (2019). A computer vision model confirmed that the movies differed in low-level features while depicting similar high-level content. Consistent with sensory exaggeration, the animated version yielded more reliable neural responses and better decoding of visual features than the CGI version throughout occipital cortex. This effect was not observed in adults nor in higher-order regions in infants and could not be explained by differential head motion or looking time. These results suggest that the developing visual system may be attuned to diagnostic features and that cartoons may (unwittingly) exploit this early neural preference.

## Introduction

Adult human vision is exquisite, supported by a highly organized and specialized visual system that efficiently processes meaningful elements of the visual world (Grill-Spector and Malach, 2004). How this system develops is an active area of research (Aslin and Bejjanki, 2024; Ayzenberg and Behrmann, 2024), with recent neuroimaging evidence of sophisticated visual system organization in human infancy (Fransson et al., 2007; Deen et al., 2017; Ellis et al., 2021; Kosakowski et al., 2022; Kubota et al., 2025; O’Doherty et al., 2026). At the same time, infancy is marked by perceptual learning and sensitive periods for the development of vision (Lewis and Maurer, 2005; Braddick and Atkinson, 2011). Thus, it is unclear whether the infant visual system is responsive to the same kinds of visual inputs as adults. Approaches that map the abstract features driving neural responses (Freiwald et al., 2009; Haxby et al., 2011; Park et al., 2022), such as by characterizing a neuron or brain region’s ‘optimal stimulus’ (Hubel and Wiesel, 1962; Tanaka et al., 1991; Ponce et al., 2019), have uncovered the computational properties of visual cortex of adults and nonhuman animals (Ratan Murty et al., 2021; Khosla et al., 2022). Such an investigation into the preferred features of the infant visual system could reveal how visual cortical areas (Kanwisher, 2010) develop specialization, with implications for the relationship between visual perception, attention, and learning in infancy (Emberson, 2019).

One possibility is that the infant visual system responds preferentially to visual features that match the natural statistics of the environment in which humans evolved and in which infants still develop (Clerkin et al., 2017). From this evolutionary perspective, the infant visual system comes equipped or rapidly gains the ability to process realistic visual information that mirrors their external environment. An early preference for realistic input may be supported by the early maturation of the infant visual system in terms of its structure (Dubois et al., 2014), connectivity (Fransson et al., 2007; Kamps et al., 2020; Kubota et al., 2025), and functional organization (Deen et al., 2017; Ellis et al., 2021; Kosakowski et al., 2022; Ellis et al., 2025).

Alternatively, infants may initially show preferential processing of ‘caricatured’ features that are both simplified and exaggerated. This sensory exaggeration hypothesis is based on the idea that such features may be beneficial for a developing organism to learn what information is important or useful in its environment. In the auditory domain, infants prefer speech characterized by higher pitch, shorter utterances, longer pauses, and exaggerated prosody (Fernald et al., 1989; Cooper and Aslin, 1990; The ManyBabies Consortium, 2020). This infant-directed speech (or ‘motherese’) facilitates language learning by enhancing vowel discrimination (Trainor and Desjardins, 2002), word segmentation (Thiessen et al., 2005), and the mapping of new words to novel objects (Ma et al., 2011), which in turn supports vocabulary growth (Weisleder and Fernald, 2013). Adults also make infant-directed actions (‘motionese’; (Brand et al., 2002)), which infants also prefer (Brand and Shallcross, 2008) and benefit from in terms of their imitation and exploration (Williamson and Brand, 2014; Meyer et al., 2023). Infant-directed speech and gesture are part of a suite of communicative behaviors that caregivers use to enhance infant perception, attention, and learning (Kosie and Lew-Williams, 2024). The infant brain may thus exhibit a preference for simplified and exaggerated visual content. Although a stimulus can be simplified without being exaggerated (e.g., a stick figure) or exaggerated without being simplified (e.g., a photorealistic visual parody), children’s animated media tends to have both properties (Wass and Smith, 2015). Indeed, infants show enhanced looking to faces during cartoons versus more realistic movies (Frank et al., 2009, 2014).

The sensory exaggeration hypothesis is inherently developmental. It predicts preferential processing of exaggerated content only early in life when useful for learning. Although infants show enhanced neural responses to infant-directed versus adult-directed speech (Zangl and Mills, 2007; Kalashnikova et al., 2018; Menn et al., 2022), adults (except mothers of preverbal infants) do not show the same effect (Matsuda et al., 2011). The same may be true for sensory exaggeration in the visual domain, or visual exaggeration may be beneficial throughout development. Indeed, adults and older children are better at perceptually discriminating caricatured faces (Rhodes et al., 1987; Chang et al., 2002) and face cells in the adult monkey brain show enhanced responses to extreme features (Leopold et al., 2006; Freiwald et al., 2009; Koyano et al., 2021). Although such studies generally focus on face caricatures, there is also evidence for privileged processing of shape caricatures in adults (Sun et al., 2024). Thus, it is unknown whether preferential processing of visual exaggeration in the brain would be specific to infants or applies throughout the lifespan.

We tested the sensory exaggeration hypothesis by examining how the infant brain responds to caricatured visual input with functional magnetic resonance imaging (fMRI). We used a movie-based design because continuous, dynamic stimuli are more engaging to infants and improve data quality and retention (Behm et al., 2026). Even still, the amount of fMRI data that can be obtained from awake infants is limited by their short attention span. We aimed to collect two versions of the same 3-minute movie sequence from each infant (order counterbalanced): a caricatured version depicted traditional hand-drawn 2-D animation (cartoon) and a realistic version depicted photorealistic 3-D computer graphics (realistic). Matching the movie sequence while manipulating the visual style allowed us to examine sensory exaggeration while controlling for many contents of the movie (e.g., presence of faces, number of camera zooms) that could influence infant attention (Tran et al., 2017; Kadooka and Franchak, 2020). We tested whether the infant visual system was better able to process the cartoon versus realistic movie with two analysis approaches. First, we used intersubject correlation (ISC) to measure ‘neural synchrony’ across infants (Hasson et al., 2004). ISC isolates the signal in fMRI activity common across participants from noise that is specific to individuals, providing a data-driven measure of the reliability of stimulus responses across the brain (Nastase et al., 2019). Second, we used multivariate pattern analysis to decode high-level visual features. We focused on decoding faces and scenes given their importance in early life and evidence of selectivity for these categories in infant visual cortex (Kosakowski et al., 2022; Kamps et al., 2025). Greater neural synchrony and better feature decoding for the cartoon versus realistic movies would support the sensory exaggeration hypothesis. We further compared these results to adults, where differences by age group would indicate that the effects are developmental in nature.

## Methods

### Participants

Data were collected from 24 infant sessions (10 female, 3 other gender; 18 unique individuals) ranging from 4 to 15 months of age (*M* = 8.38, *SD* = 3.44 months). Six infants completed more than one usable session (*M* = 1.66 sessions, range: 1 to 3 sessions), with an average of 2.31 months between consecutive sessions (range: 1.20 to 3.60 months). These sessions were treated as independent, consistent with our prior work (Ellis et al., 2020b; Yates et al., 2022; Ellis et al., 2025). The final sample does not include data from infants who did not watch either movie in full (N = 5), only saw one movie in full (N = 9), had excessive head motion (>3-mm framewise displacement) for more than 50% of one or both movies (N = 3), or did not look at the screen for more than 50% of a movie (N = 1). For comparison, we collected movie-watching data from 12 adults (7 female) aged 19 to 29 years (*M* = 21.42, *SD* = 3.40 years), all of whom had usable data.

Infants were recruited through the Yale Baby School, an outreach initiative to families who give birth at the Yale New Haven Hospital. Adults were recruited from the New Haven, Connecticut area. This study was approved by the Institutional Review Board (IRB) at Yale University. All adults provided informed consent, and parents provided informed consent on behalf of their infant.

## Materials

We created two silent 3-minute movie clips from the 1994 and 2019 versions of the film, The Lion King. These clips depicted the same opening sequence of the movie, with multiple animal groups moving through the savanna to a rock formation where a baby lion is introduced to the animal kingdom (Figure 1A). Importantly, the 1994 version was hand-drawn 2D traditional animation (referred to as the ‘cartoon’ condition) whereas the 2019 version was photorealistic 3D computer-generated animation (referred to as the ‘realistic’ condition). Movies were re-cut in iMovie to align the timing and match the semantic content between versions as precisely as possible on a frame-by-frame basis. The auditory track was removed from both clips.

**Figure 1.**
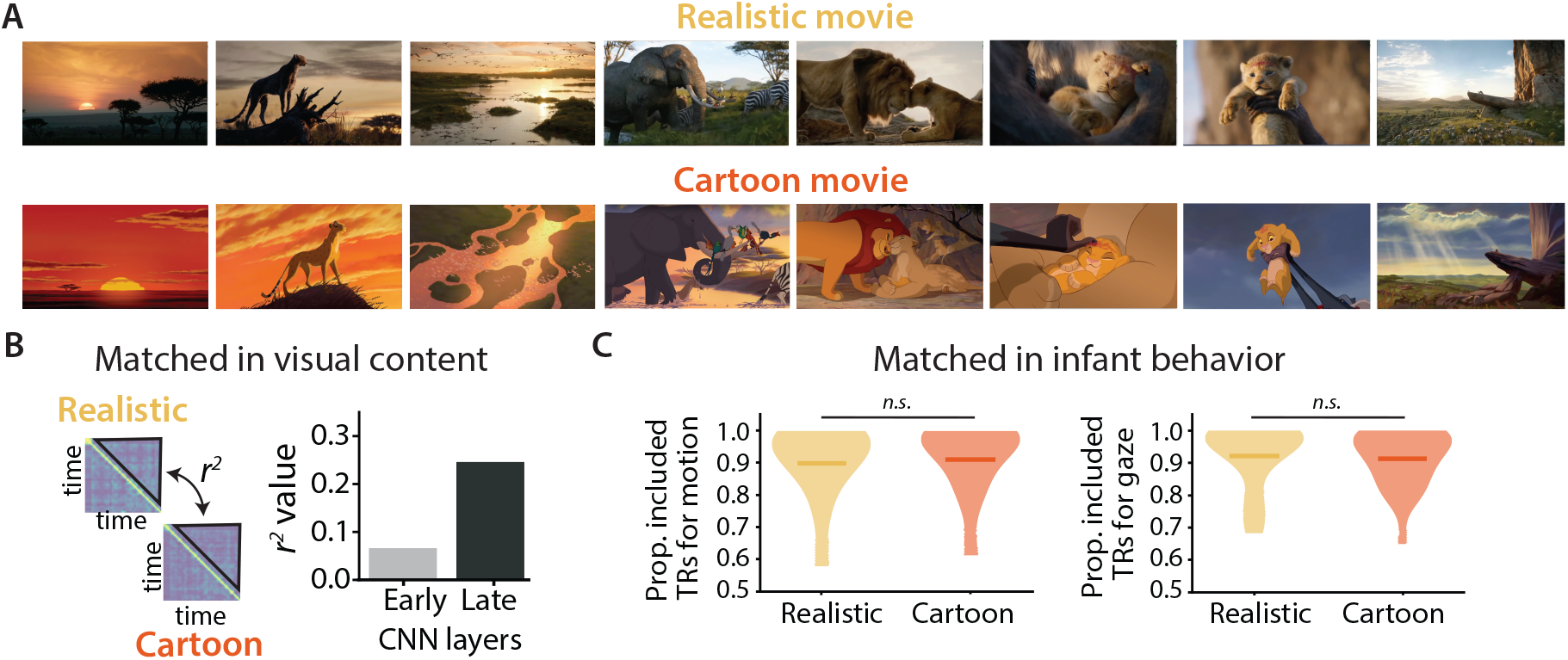
Awake infant fMRI movie task. (A) Timelocked example frames from the realistic (top) and cartoon (bottom) movies shown to infants and adults. (B) The movies differed in low-level style but were better matched in high-level content, leading to dissimilar representations in early layers of the VGG-19 computer vision model (gray, max pooling layer 2) and more similar representations in late model layers (dark gray, max pooling layer 5). See Figure S1 for all layers. (C) Infant behavior did not differ between the realistic (yellow) and cartoon (orange) movies in terms of the proportion of timepoints retained after exclusions for excessive head motion and inattentive gaze. Violin plot of distribution across participants with the mean indicated by a saturated horizontal line. n.s. = not significant.

The code used to show the movies on the experimental display is available at https://github.com/ntblab/experiment_menu/tree/CartoonLive. The code used to perform the data analyses is available at https://github.com/ntblab/infant_neuropipe/tree/CartoonLive. Raw and preprocessed functional data and anatomical images will be made available upon publication. The movies shown to infants and adults are copyrighted but available upon request.

### Data acquisition

Data were collected at the Brain Imaging Center (BIC) in the Faculty of Arts and Sciences at Yale University. We used a Siemens Prisma (3 Tesla) MRI with the bottom half of the 20-channel Siemens head coil for both infant and adult participants. Functional images were acquired with a whole-brain T2* gradient-echo echoplanar imaging sequence (TR = 2 s, TE = 30 ms, flip angle = 71, matrix = 64×64, slices = 34, resolution = 3 mm iso, interleaved slice acquisition). Anatomical images were acquired with a T1 PETRA sequence for infants (TR1 = 3.32 ms, TR2 = 2250 ms, TE = 0.07 ms, flip angle = 6, matrix = 320×320, slices = 320, resolution = 0.94 mm iso, radial lines = 30,000) and a T1 MPRAGE sequence for adults, with the top of the head coil attached (TR = 2400 ms, TE = 2.41 ms, TI = 1000 ms, flip angle = 8, iPAT = 2, slices = 176, matrix = 256×256, resolution = 1.0 mm iso).

### Procedure

We collected awake fMRI data from infants and adults using a previously validated procedure (Ellis et al., 2020b). Before their first scan, infants and their parents met with researchers for an orientation session either in-person or online, in accordance with COVID-19 policies. The fMRI scan session was then scheduled for a time when the parent thought the infant was likely to be happy and awake. Parents and infants were screened carefully for metal in advance and on the day of the scan. Experimenters applied three layers of hearing protection on the infants: silicon inner ear putty, over-ear adhesive covers, and passive headphones. Infants were placed on a vacuum pillow that comfortably reduced their movements and parents were encouraged to bring metal-free comfort items (e.g., blankets, pacifiers) for their infant. The top of the head coil was not placed over the infant in order to maintain comfort. Stimuli were projected onto the ceiling of the scanner bore above the infant’s face using a mirror projection system. We recorded video of the infant’s face (MRC high-resolution camera) during the session for offline gaze coding. Adult participants underwent the same procedure with the following exceptions: they did not attend an orientation session, hearing protection was only two layers (earplugs and Optoacoustics noise-canceling headphones), they were not on a vacuum pillow, and they were not given comfort items. Additional tasks, not described here, were sometimes run during infant and adult scanning sessions.

Experimental stimuli were shown using Psychtoolbox (Kleiner et al., 2007) for MATLAB. The projected visual stimuli extended 45.0 degrees of visual angle wide by 25.5 degrees tall. Half of the infants (N = 12) saw the cartoon version of the movie first and the other half (N = 12) saw the realistic version of the movie first. All but two participants saw each movie only once in a given session. One participant (4.0 months) saw the cartoon version twice because the first acquisition had movement-related aliasing in the functional image; we only analyzed data from the second viewing without aliasing. Another participant (15.0 months) saw part of the realistic version before it was stopped and restarted because of technical issues with the camera for gaze monitoring. In all but two sessions, the two movies were shown back-to-back in the same functional run; in the other cases, there was a short delay between the movies (719.69 and 831.27 s) for another task, anatomical scan, and/or break. In adults, half of the participants (N = 6) saw the cartoon version first and the other half (N = 6) saw the realistic version first. Adults watched other movies, not described here, and completed a fixation-cross resting-state scan in a counterbalanced order (Yates et al., 2023a). In all adult sessions, movies were presented in separate functional runs. In some adult sessions (N = 3), the two movies were shown in back-to-back runs; otherwise, they were separated by a short delay (mean = 438.38 s, range = 406.70 to 465.62) for other functional scans.

### Gaze coding

Infant gaze was coded offline by 2-3 coders (*M* = 2.13 *SD* = 0.33). Coders determined whether the participant’s eyes were on-screen, off-screen (i.e., closed, blinking, or looking away), or undetected (i.e., out of the camera’s field of view). Coders were highly reliable, reporting the same response code on an average of 96.65% (*SD* = 2.22%; range across participants = 91.90 to 99.68%) of frames for the cartoon movie and 95.39% (*SD* = 4.14%; range across participants = 86.16 to 99.73%) of frames for the realistic movie. The modal response across coders from a moving window of five frames was used to determine the final response for the frame centered in that window, with the response from the previous frame used in the case of ties. Frames were then pooled within TRs, and we analysed the average proportion of TRs in which infants were deemed to be looking on-screen. Gaze data were coded by one coder for two of the adults who were noted to be sleepy during data acquisition. One of these adults maintained fixation for 98.9% of frames for both movies, while the other adult was attentive during their first movie viewing (realistic movie, 95.6% of frames looking on-screen) and falling asleep during their second movie viewing (cartoon movie, 58.2% of frames looking on-screen). Because this adult was looking on-screen for more than half of the movie (same threshold used in infants), they were included in the analysis but their off-screen timepoints were excised.

### Preprocessing

Data were preprocessed using a modified FSL FEAT pipeline designed for infant fMRI (Ellis et al., 2020b). If infants participated in other tasks during the same functional run (*N* = 13), the movie data were cleaved to create a pseudo-run. Three burn-in volumes were discarded from the beginning of each run/pseudo-run.

Motion correction was applied using the centroid volume as the reference, determined by calculating the Euclidean distance between all volumes and choosing the volume that minimized the distance to all other volumes. Slices in each volume were realigned using slice-timing correction. Timepoints with greater than 3 mm of translational motion were interpolated from surrounding timepoints so as not to bias linear detrending then excluded from later analyses. All adult timepoints were included for both movies, as no timepoints exceeded the motion threshold.

Timepoints during which eyes were closed for more than half of the 48 frames in the volume (frame rate of 24 frames-per-second for 2-s TR) were excluded from subsequent analyses. The signal-to-fluctuating-noise ratio (SFNR) (Friedman and Glover, 2006) was calculated for each voxel and thresholded to form a mask of brain versus non-brain voxels. Data were spatially smoothed with a Gaussian kernel (5-mm FWHM) and linearly detrended in time. AFNI’s (https://afni.nimh.nih.gov) despiking algorithm was used to attenuate aberrant timepoints within voxels. After removing excess burn-out TRs, functional data were *z*-scored within run.

The centroid functional volume was registered to the PETRA anatomical image. Initial alignment was performed using FLIRT with 6 degrees of freedom (DOF) and a normalized mutual information cost function. This automatic registration was manually inspected and then corrected if necessary using mrAlign from mrTools (Gardner lab). To compare across participants, infant PETRA anatomical images were aligned to an age-specific infant MNI template (Fonov et al., 2009) using an initial linear alignment with 12 DOF, followed by non-linear warping using diffeomorphic symmetric normalization (ANTs; (Avants et al., 2011)). Then, we used a predefined transformation (12 DOF) to linearly align the age-specific infant MNI template to the adult MNI standard (1-mm voxel resolution). For adults, we used the same alignment procedure, except that the adult anatomical images (MPRAGE) were directly aligned to the adult MNI standard. For all analyses, we only considered voxels in adult MNI standard space included in the intersection of all infant and adult brain masks. Analyses were performed in this standard space at the native voxel resolution (3 mm).

### Regions of interest

We performed analyses over the whole brain and in regions of interest (ROIs). For visual ROIs, we used a probabilistic atlas of topographic visual areas constructed based on retinotopic mapping in adults (Wang et al., 2015). We used ROIs at the maximum probability level and combined them into three larger ROIs: early visual (comprising V1v and V1d), lateral occipital (comprising LO1 and LO2), and ventral occipital (comprising VO1 and VO2). For comparison, we also chose two ROIs that we did not expect to be modulated by visual style: the precuneus, a higher-order brain region involved in narrative comprehension, and early auditory cortex, given that the movie as presented was silent; these ROIs were created using the Harvard-Oxford probabilistic atlas (Jenkinson et al., 2012) (0% probability threshold).

For face and scene decoding, we focused on the ventral occipital cortex ROI, as it encompasses regions important for high-level visual category processing in adults (Kanwisher, 2010). We additionally examined more specific category-selective regions: the bilateral fusiform face area (FFA) and bilateral parahippocampal place area (PPA). We defined FFA and PPA ROIs by creating 10-mm radius spheres around the peaks in activation in meta-analytic maps from Neurosynth (Yarkoni et al., 2011), as previously used in our research (Yates et al., 2023b). In exploratory analyses of MR-based eye-tracking (Frey et al., 2021; Nau et al., 2025), we manually defined left and right eyeball ROIs in each infant’s anatomical image and retained eyeball voxels that were common across all infant participants. All ROIs were transformed into the fMRI voxel resolution (3 mm).

### Movie labeling

For decoding analyses, one of the authors (T.S.Y.) manually coded each movie separately for two regressors: face presence (present/absent) and scene distance (close/medium/far). These features were coded for each 2-s segment of the movie, corresponding to a TR in the fMRI data shifted 4 s to account for hemodynamic lag. For the face regressor, a segment was coded as ‘1’ if at least one (animal) face was present and identifiable (eyes and/or mouth visible); otherwise, it was coded as ‘0’. For the scene regressor, a segment was coded as a ‘0’ if the camera angle was close-up, ‘1’ if it was at a medium distance, and ‘2’ if it was at a far distance, following the coding scheme used in a previous fMRI movie-watching dataset in adults (Chen et al., 2017).

### Computer vision model

We first verified that the two movie versions with different low-level visual styles conveyed similar high-level semantic content. Movie frames were analyzed using VGG-19 (Simonyan and Zisserman, 2014), a computer vision model with a deep convolutional network architecture trained to perform image classification. Each frame of the cartoon or realistic movie was fed into a pretrained VGG-19 architecture obtained through Keras (Chollet et al., 2015) and Pytorch (Paszke et al., 2019). For each max pooling and fully connected layer of the network, the layer unit values across frames were convolved with the hemodynamic response function (Ellis et al., 2020a), resulting in a layer unit by TR matrix. These matrices were then transformed into TR-by-TR correlation matrices by taking the Pearson correlation between layer unit values for each pair of TRs. To compare model representations for the two movies, we calculated the second-order Pearson correlation between the upper triangles of these TR-by-TR matrices for each layer; the diagonal and first 4 off-diagonals were excluded (similarity of a timepoint to itself or to the next 4 timepoints, respectively) to account for autocorrelation. We examined all layers of the network, with the logic that lower layers capture low-level visual features and higher layers capture high-level semantic content.

### Visual complexity measures

Movie frames were additionally analyzed for the complexity of visual features. We used composite measures of three features: shape, color, and motion. Complexity measures were calculated every 500 ms using OpenCV (Bradski, 2000) and aggregated into 2-s epochs, corresponding to TRs. Shape complexity comprised a weighted sum of a frame’s multi-scale Canny edge density (Canny, 1983), orientation entropy (Sobel et al., 2022), and corner density (Shi and Tomasi, 1994). Color complexity was calculated as a weighted sum of the Shannon entropy across pixel values in AB color space (hue/saturation values) and the standard deviation in luminance from that frame, with luminance values normalized across the movie. Motion complexity was calculated as the optical flow magnitude taking into account the speed, direction, and coherent motion between the previously analyzed frame and the current frame (Farnebäck, 2003), with motion magnitude normalized across the movie.

### Intersubject correlation analysis

We assessed neural synchrony across participants for both movie types (cartoon, realistic) using ISC (Hasson et al., 2004; Nastase et al., 2019). For each voxel, we calculated the Pearson correlation of the timecourse of BOLD activity between a single held-out participant and the average timecourse of all other participants in the same age group (infant or adult); the resulting voxelwise map of correlation coefficients was Fisher-transformed. We iterated such that each participant was held-out once and averaged the maps across iterations.

Whole-brain maps of ISC were generated using non-parametric statistical tests of the reliability of the effect above chance across participants at the group level (Winkler et al., 2014). For each movie type and age group, we used FSL’s randomise tool to conduct a randomization test over the Fisher-transformed voxelwise ISC maps from all participants. Specifically, the sign of each voxel’s data was randomly flipped on each of 1,000 iterations under the null hypothesis of no difference from chance, where the *p*-value was the proportion of permutations with a more extreme value than the actual data. We used threshold-free cluster enhancement (TFCE) for multiple comparisons correction (Smith and Nichols, 2009). To compare across movie types, a similar randomization test was performed after first subtracting the voxelwise ISC maps for the cartoon minus realistic movies within participant.

For the ROI analysis, Fisher-transformed ISC values were averaged across voxels within the region for each held-out participant. Statistical significance across participants was determined with bootstrap resampling (Efron and Tibshirani, 1986). We randomly sampled participants with replacement 1,000 times, on each iteration forming a new sample of the same size as the original group and averaging their ISC values to generate a sampling distribution across iterations. The *p*-value was calculated as the proportion of re-sampling iterations in which the group average had the opposite sign as the original effect, doubled for a two-tailed test. To compare across movie types, we subtracted the ISC values for the cartoon minus realistic movies within participant, and then conducted bootstrap resampling of the differences across participants as above.

### Multivariate pattern analysis

We decoded face presence and scene distance from BOLD activity using multivariate pattern analysis (Norman et al., 2006). We adopted an across-participant correlation-based classification method that has been used previously in infant fNIRS (Emberson et al., 2017). This method can handle small amounts of training data without overfitting because it does not learn weights for each voxel and instead treats them equally. Namely, for each movie type and classifier (e.g., face presence) we built a template representation for each label (e.g., present, absent) by averaging the pattern of voxel activity for all of the timepoints with that label in all but one held-out participant. Then, for each timepoint in the held-out participant’s data, we calculated the correlation of the activity pattern with the template representation (from the other participants) for each label. Classification accuracy was defined as the proportion of timepoints in which the correlation was higher for the template corresponding to the correct label (e.g., face present template for a timepoint when a face was present) than the incorrect template (e.g., face absent template when face present). Chance accuracy was 50% for face presence and 33% for scene distance (i.e., because there are three possible labels for that classifier).

Decoding was attempted both in a whole-brain searchlight analysis (Kriegeskorte et al., 2006; Kumar et al., 2021) and inside ROIs. For the searchlight analysis, we defined a spherical mask around each brain voxel (radius = 3 voxels) and extracted vectorized activity patterns from all voxels in the mask; classification accuracy was assigned to the center voxel of each searchlight. Whole-brain statistical maps were created by subtracting chance accuracy from each voxel’s classification accuracy for each participant, using FSL’s randomise tool to conduct a randomization test, then correcting for multiple comparisons with TFCE. To assess differences across movie types, a similar randomization test was performed after first subtracting the voxelwise classification accuracy maps for the cartoon minus realistic movies within participant.

For the ROI analysis, statistical significance for each movie type was determined using the bootstrap resampling approach described above, this time with the *p*-value calculated as the proportion of resampling iterations in which the group average classification accuracy was below the chance level, doubled for a two-tailed test. To compare across movies, we subtracted the classification accuracy values for cartoon minus realistic movies for each participant and then conducted bootstrap resampling of the difference as above.

### Supplementary movie experiments

This study was inspired by serendipitous observations made while collecting other infant movie-watching fMRI datasets in prior sessions. These datasets include one study with 24 infants aged 3.6 to 12.7 months (*M* = 7.43, *SD* = 2.70; 12 female) watching a silent cartoon (“Aeronaut”), as described in (Yates et al., 2022, 2023a); the movie extended 45.0° wide by 25.5° high and lasted 3 minutes. Another dataset had 22 infants aged 3.3 to 32.0 months (*M* = 13.67, *SD* = 7.86; 13 female) watching four silent, live-action video clips concatenated in the same order (“Child Play”), as described in (Ellis et al., 2025); the movie extended 40.8° wide by 25.5° high and lasted 6.8 minutes. Both datasets were collected at the Brain Imaging Center at Yale University and preprocessed using the pipeline as described above. All parents provided informed consent on behalf of their infants for these prior studies.

We report supplementary results for the comparison of Aeronaut to Child Play, but do not consider these primary data because the movies were different in multiple ways beyond the key contrast of cartoon versus realistic, most critically in the contents, size, and duration of the movies. Thus, any differences must be interpreted with caution, and served only as inspiration for selecting more controlled stimuli in the current study. That said, the results generally agree and thus provide additional support for our conclusions.

## Results

### Matched high-level and different low-level visual features in realistic and cartoon movies

We first assessed whether the two movies (cartoon and realistic) were matched as intended for content despite different visual styles. Namely, we hypothesized that they would have similar high-level, abstract features despite dissimilar low-level, perceptual features. We used a convolutional neural network (CNN) model, VGG-19 (Simonyan and Zisserman, 2014), to extract visual features from the movie frames. CNNs can predict neural responses across the visual processing hierarchy in monkeys and humans (Yamins et al., 2014; Yamins and DiCarlo, 2016), with earlier layers of the network corresponding to earlier visual areas (i.e., V1) and later layers of the network corresponding to later visual areas (i.e., inferior temporal cortex).

We predicted that the two movies would be represented more similarly in later than earlier layers of VGG-19. Indeed, the variance shared between model representations of the movies increased from earlier layers of the network (max block pooling layer 2: *r*^2^ = 0.07) to later layers of the network (max block pooling layer 5: *r*^2^ = 0.26; Figure 1B), as did the magnitude of the relationship (max block pooling layer 2: *β* = 0.23; max block pooling layer 5: *β* = 0.68). This increasing similarity was not a general property of the model architecture, as more variance was shared between the realistic and cartoon versions of the same movie compared to an unrelated movie (Figure S1).

As a further manipulation check, we analyzed the movies using composite visual complexity measures of color, shape, and motion derived from computer vision algorithms (Bradski, 2000). Color complexity (operationalized as a weighted sum of hue entropy and relative luminance variance) was lower for the realistic (mean over time, *M* = 0.497, *SD* = 0.105) than cartoon movie (*M* = 0.547, *SD* = 0.120; difference: *M* = −0.050, 95% confidence interval, CI = [−0.083 to −0.016], bootstrap *p* <0.001), reflecting a lower spread of color values (Figure S2A). In contrast, shape complexity (operationalized as a weighted sum of Canny edge density, corner density, and orientation entropy) was higher for the realistic (*M* = 0.522, *SD* = 0.052) than cartoon movie (*M* = 0.388, *SD* = 0.089; difference: *M* = 0.134, CI = [0.113 to 0.155], *p* <0.001), reflecting greater edge and corner density. Motion complexity (operationalized as optical flow magnitude between frames taking into account speed, direction, and coherent motion) was not significantly different between the realistic (*M* = 0.670, *SD* = 0.106) and cartoon movie (*M* = 0.689, *SD* = 0.109; difference: *M* = −0.020, CI = [−0.052 to 0.010], *p* = 0.196). Altogether, these computer vision analyses validate that the movie versions manipulated visual style in terms of color and shape, while maintaining similar abstract visual content and motion.

### Comparable infant behavior during realistic and cartoon movies

We next assessed data quality and infant attentiveness for the two movies (Figure 1C). There was no difference in the proportion of fMRI timepoints (TRs) retained after exclusion for head motion between the cartoon (mean over participants, *M* = 91.0%, *SD* = 11.3%) and realistic movie (*M* = 89.8%, *SD* = 12.1%; difference: *M* = 1.2%, CI = [−2.8% to 5.3%], bootstrap *p* = 0.624). Relatedly, there was no difference in the amount of head motion as measured by median framewise displacement between the cartoon (*M* = 0.28 mm, *SD* = 0.22 mm) and realistic movie (*M* = 0.28 mm, *SD* = 0.23 mm; difference: *M* = −0.003 mm, CI = [−0.06 mm to 0.06 mm], *p* = 0.942).

Infants were also equally attentive to the two movies. There was no difference in the proportion of TRs retained after exclusion for gaze off-screen or eyes closed between the cartoon (*M* = 91.3%, *SD* = 8.9%) and realistic movie (*M* = 92.2%, *SD* = 10.0%; difference: *M* = −0.9%, CI = [−4.5% to 2.8%], *p* = 0.634). Furthermore, spatial ISC in the eyeballs, which has been used as an MRI-based eyetracking signal in adults (Frey et al., 2021; Nau et al., 2025), was equally reliable for the realistic and cartoon movies and similar between movies (Figure S3). These results suggest that neural differences between the two movies cannot easily be attributed to differences in head motion, attention, or data quality more generally.

### Enhanced stimulus-driven neural synchrony for cartoon movies in infant visual cortex

To investigate the reliability of neural responses to the movies across infants, we calculated leave-one-out intersubject correlation (ISC). For every voxel, the timecourse of BOLD activity from one participant was correlated with the average timecourse from all other participants (Hasson et al., 2004); this was repeated 24 times, each time leaving out a different participant. In addition to examining the whole brain, we also calculated ISC within five ROIs: three visual regions (early visual, lateral occipital, ventral occipital cortices), a higher-order multimodal region associated with narrative processing (Baldassano et al., 2017) (precuneus), and a control region for another sensory modality (early auditory).

Across the whole brain, ISC was most robust and survived multiple comparisons correction in occipital and frontal cortices for both the realistic and cartoon movies (Figure 2A). The magnitude and extent of ISC was notably greater for the cartoon movie in visual cortex and orbitofrontal cortex (Figure S4A); the realistic movie led to reliable but weak ISC in cingulate cortex and medial prefrontal cortex.

**Figure 2.**
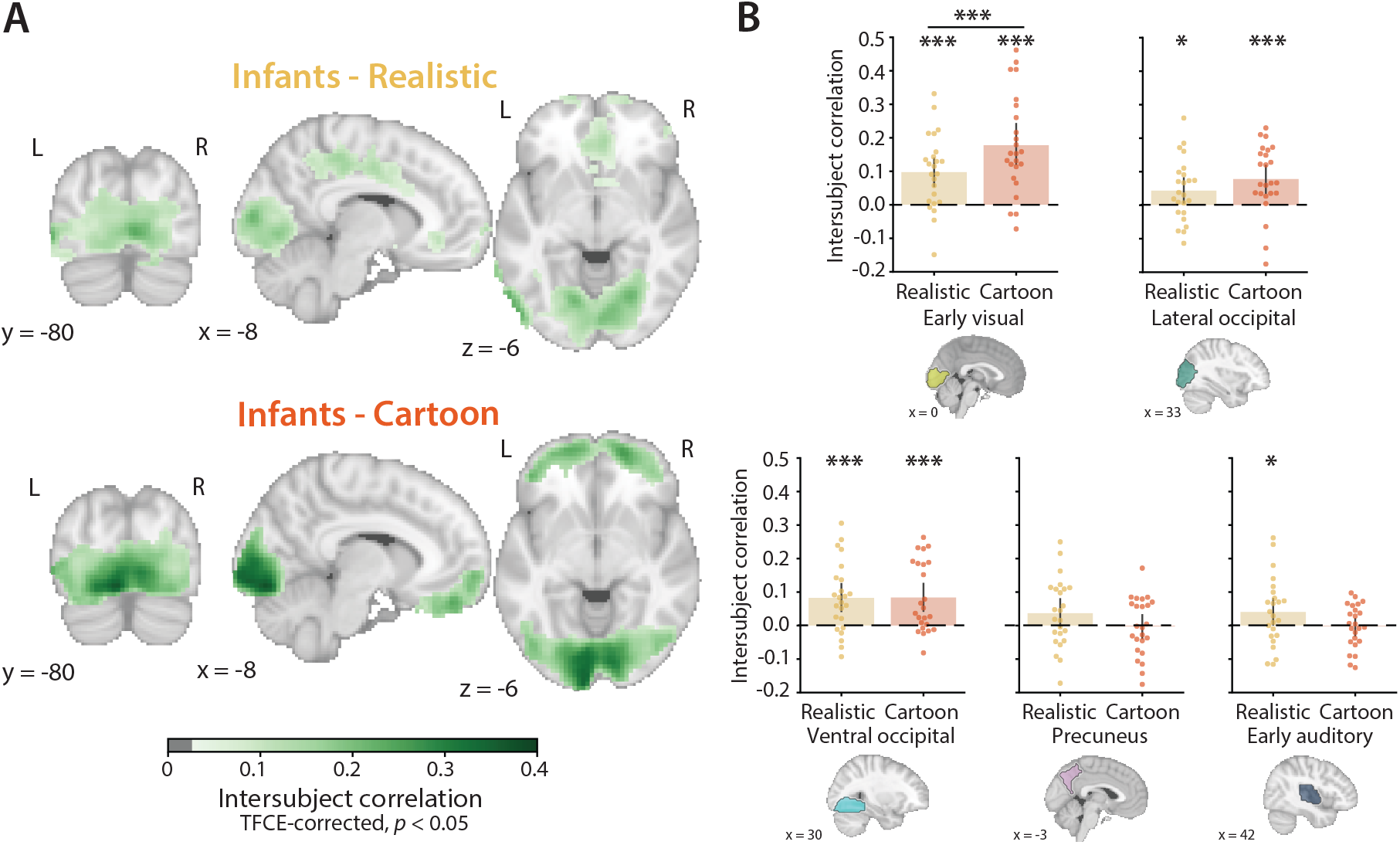
Neural synchrony for realistic and cartoon movies in infants. (A) Voxel-wise ISC thresholded at *p* <.05, corrected with TFCE. (B) ISC was significant in all three visual ROIs (early visual, lateral occipital, ventral occipital) for both the realistic and cartoon movies. Importantly, ISC was significantly higher for the cartoon than realistic movie in early visual cortex. Dots represent individual participants and error bars represent 95% CIs of the mean from bootstrap resampling. *** *p* < 0.001, * *p* < 0.05.

The ROI analysis was compatible with the whole-brain result from occipital cortex (Figure 2B). In early visual cortex, ISC was significantly greater than chance for both realistic (*M* = 0.098, CI = [0.055 to 0.141], bootstrap *p* < 0.001) and cartoon movies (*M* = 0.178, CI = [0.123 to 0.238], *p* < 0.001), and was significantly stronger for the cartoon movie (*M* = 0.080, CI = [0.024 to 0.142], *p* < 0.001). In lateral occipital cortex, ISC was significant for both realistic (*M* = 0.044, CI = [0.010 to 0.082], *p* = 0.014) and cartoon movies (*M* = 0.079, CI = [0.036 to 0.118], *p* < 0.001); although ISC was numerically stronger for the cartoon movie, this difference was not reliable (*M* = 0.035, CI = [−0.007 to 0.080], *p* = 0.110). In ventral occipital cortex, ISC was again significant for both realistic (*M* = 0.083, CI = [0.044 to 0.127], *p* < 0.001) and cartoon movies (*M* = 0.084, CI = [0.041 to 0.127], *p* < 0.001), which did not differ (*M* = 0.001, CI = [−0.051 to 0.055], *p* = 0.970). In precuneus, ISC was not significant for either realistic (*M* = 0.037, CI = [−0.004 to 0.077], *p* = 0.092) or cartoon movies (*M* = −0.003, CI = [−0.038 to 0.031], *p* = 0.822), with a numerical trend toward stronger ISC for the realistic movie (*M* diff = −0.040, CI = [−0.084 to 0.004], *p* = 0.082). In early auditory cortex, ISC was unexpectedly significant for the realistic movie (*M* = 0.041, CI = [0.001 to 0.079], *p* = 0.048) but not the cartoon movie (*M* = −0.002, CI = [−0.029 to 0.024], *p* = 0.826), with a numerical trend toward stronger ISC for the realistic movie (*M* diff = −0.043, CI = [−0.087 to 0.005], *p* = 0.062).

### Enhanced feature decoding in the infant brain during cartoon movie

One prediction of the sensory exaggeration hypothesis is that preferential processing of exaggerated features early in life enhances processing of diagnostic information. To test this prediction, we used multivariate pattern analysis to decode high-level visual features in each movie across participants, focusing on face and scene processing (Figure 3A).

**Figure 3.**
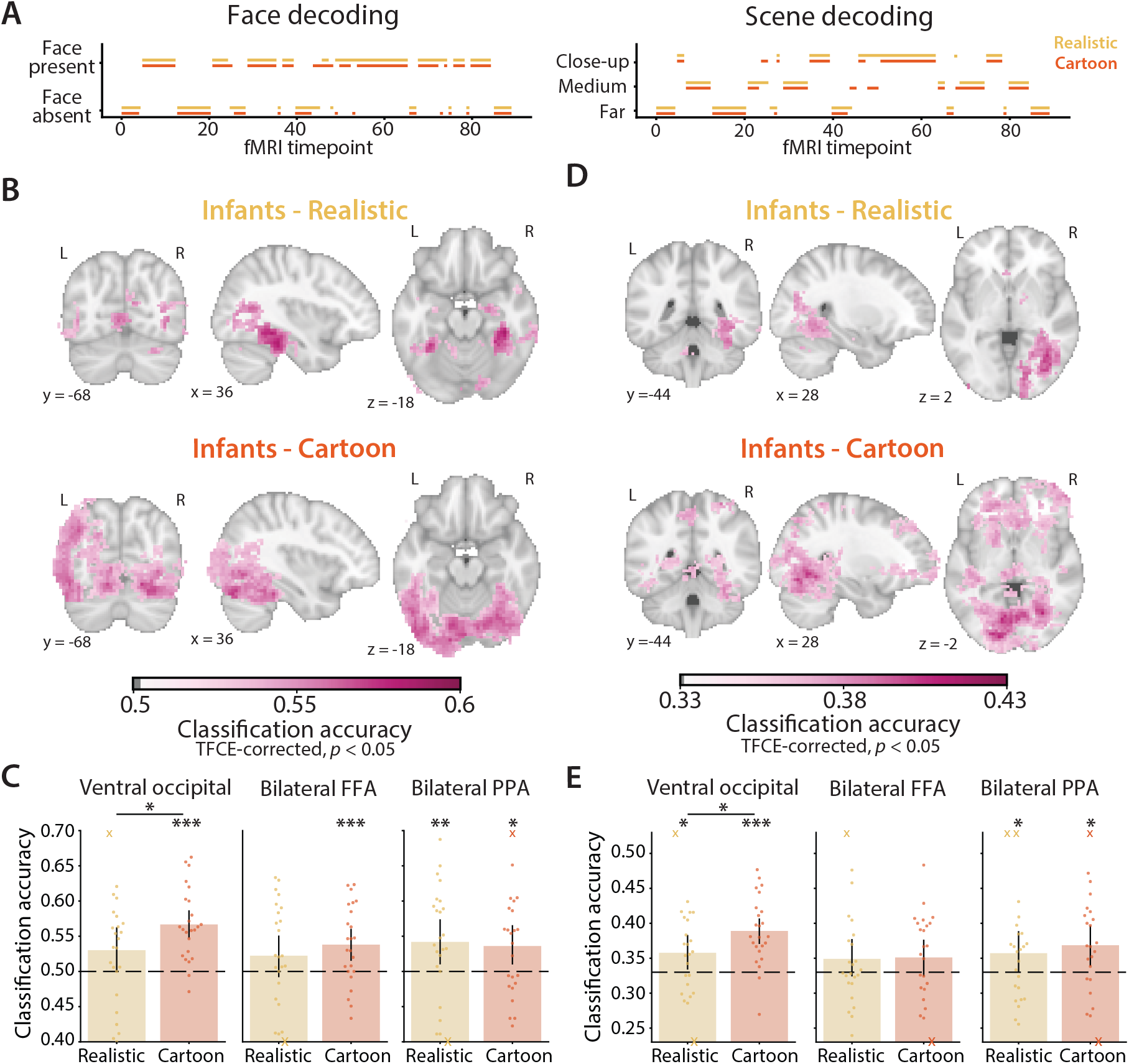
Neural decoding from realistic and cartoon movies in infants. (A) Timecourse of face and scene features used in the multivariate pattern analyses for the realistic (yellow) and cartoon (orange) movies. (B) Whole-brain searchlight results and (C) ROI results for the decoding of face presence vs. absence. (D) Whole-brain searchlight results and (E) ROI results for the decoding of close-up, medium, or far-away scene distance. In the searchlight maps, voxels significant at *p* <0.05 (corrected) are colored by the average classification accuracy across participants. For the ROIs, dots represent the classification accuracy for individual participants and error bars represent 95% CIs of the mean from bootstrap resampling. Note that (C) and (E) have different minimum and maximum values on the y-axis to highlight the range of individual values. Points outside of plotting range are marked with Xs on the positive or negative end of the y-axis. *** *p* < 0.001, ** *p* < 0.01, * *p* < 0.05. FFA = fusiform face area, PPA = parahippocampal place area.

Across the whole brain, face presence versus absence could be classified in occipital and temporal cortices for both the realistic and cartoon movies (Figure 3B). Decoding was significant across a broader swath of visual cortex for the cartoon movie, and greater than the realistic movie in regions of the occipital cortex (Figure S4B). In the ROI analysis (Figure 3C), there was significant classification of face presence versus absence for the cartoon movie in ventral occipital cortex (*M* = 0.567, CI = [0.547 to 0.587], bootstrap *p* vs. 50% chance < 0.001), bilateral FFA (*M* = 0.538, CI = [0.515 to 0.561], *p* <0.001), and bilateral PPA (*M* = 0.536, CI = [0.510 to 0.561], *p* = 0.010). For the realistic movie, face classification was only significant in bilateral PPA (*M* = 0.542, CI = [0.510 to 0.574], *p* = 0.008); there was a numerical trend in ventral occipital cortex (*M* = 0.530, CI = [0.500 to 0.563], *p* = 0.054) and no effect in bilateral FFA (*M* = 0.523, CI = [0.492 to 0.551], *p* = 0.148). Classification accuracy was significantly higher for the cartoon than realistic movie in ventral occipital cortex (*M* = 0.037, CI = [0.004 to 0.072], *p* = 0.020), but not bilateral FFA (*M* = 0.016, CI = [−0.018 to 0.049], *p* = 0.386), or bilateral PPA (*M* = −0.006, CI = [−0.043 to 0.051], *p* = 0.792). In sum, decoding of face presence versus absence was significant across visual regions for the cartoon movie, and higher than the realistic movie in ventral occipital cortex.

This same general pattern was found for scene distance decoding (Figure 3D). Although scene distance could be decoded from the realistic movie in a cluster spanning right occipital and temporal cortices, decoding was stronger and broader for the cartoon movie, particularly in occipital regions (Figure S4C). In the ROI analysis (Figure 3E), there was significant classification of scene distance for the cartoon movie in ventral occipital cortex (*M* = 0.389, CI = [0.370 to 0.407], *p* vs. 33% chance < 0.001) and bilateral PPA (*M* = 0.369, CI = [0.339 to 0.396], *p* = 0.010), but not bilateral FFA (*M* = 0.351, CI = [0.324 to 0.377], *p* = 0.122). For the realistic movie, ventral occipital cortex (*M* = 0.358, CI = [0.334 to 0.383], *p* = 0.028) and bilateral PPA (*M* = 0.357, CI = [0.330 to 0.389], *p* = 0.046) showed significant decoding but not bilateral FFA (*M* = 0.349, CI = [0.324 to 0.377], *p* = 0.154). Classification accuracy was again significantly higher for the cartoon versus the realistic movie in ventral occipital cortex (*M* = 0.031, CI = [0.006 to 0.057], *p* = 0.014), but not bilateral FFA (*M* = 0.002, CI = [−0.031 to 0.031], *p* = 0.900) or bilateral PPA (*M* = 0.011, CI = [−0.037 to 0.054], *p* = 0.594).

### No advantage for cartoons over realistic movies in adults

Our results thus far could reflect a general bias for the processing of cartoon versus realistic movies in the human brain, rather than a developmental preference. We therefore repeated the same analyses in a sample of young adults (N = 12) collected using the same protocol.

Adults showed robust and highly similar ISC for both realistic and cartoon movies in the whole-brain analysis (Figure S5A). In the ROI analysis (Figure 4A), ISC was significantly greater than chance in all ROIs for the realistic movie (early visual: *M* = 0.394, CI = [0.343 to 0.444], bootstrap *p* < 0.001; lateral occipital: *M* = 0.274, CI = [0.233 to 0.314], *p* < 0.001; ventral occipital: *M* = 0.308, CI = [0.262 to 0.360], *p* < 0.001; precuneus: *M* = 0.084, CI = [0.052 to 0.117], *p* < 0.001; early auditory: *M* = 0.038, CI = [0.009 to 0.070], *p* = 0.004) and in all but early auditory cortex for the cartoon movie (early visual: *M* = 0.401, CI = [0.326 to 0.472], *p* < 0.001; lateral occipital: *M* = 0.278, CI = [0.223 to 0.332], *p* < 0.001; ventral occipital: *M* = 0.333, CI = [0.266 to 0.392], *p* < 0.001; precuneus: *M* = 0.088, CI = [0.020 to 0.148], *p* = 0.018; early auditory: *M* = −0.006, CI = [−0.029 to 0.012], *p* = 0.580). Notably different from infants, the cartoon movie did not yield higher ISC than the realistic movie in early visual cortex (*M* = 0.007, CI = [−0.043 to 0.058], *p* = 0.800). There was also no difference in the other ROIs (lateral occipital: *M* = 0.004, CI = [−0.043 to 0.053], *p* = 0.894; ventral occipital: *M* = 0.025, CI = [−0.022 to 0.072], *p* = 0.314; precuneus: *M* = 0.004, CI = [−0.059 to 0.056], *p* = 0.880), with the exception of early auditory cortex, which showed higher ISC for the realistic than cartoon movie (*M* = 0.044, CI = [−0.006 to 0.086], *p* = 0.018).

**Figure 4.**
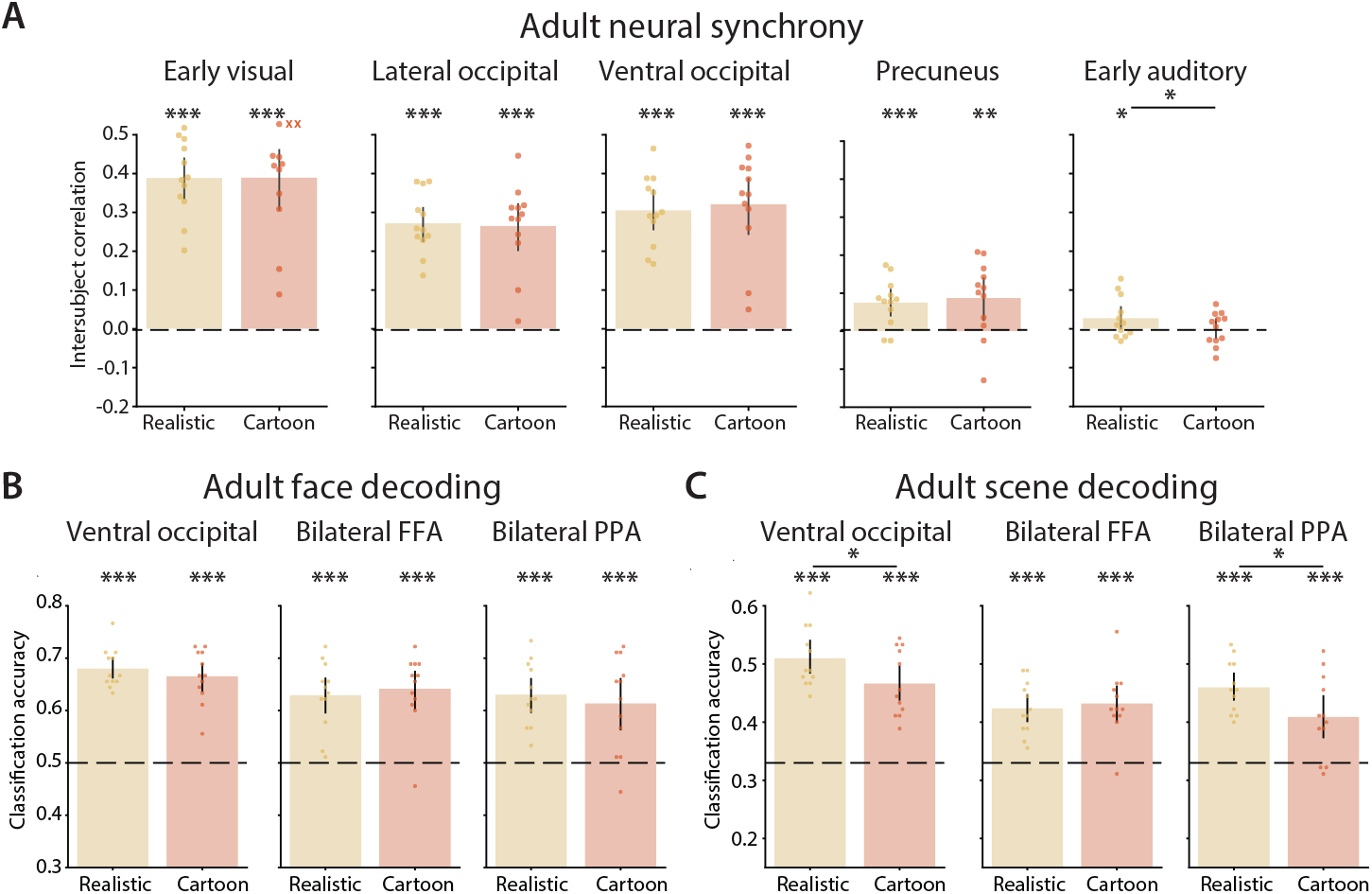
Parallel analyses in adults. (A) ROI results for ISC across adults. (B) ROI results for the decoding of face presence and (C) scene distance. Dots represent individual participants and error bars represent 95% CIs of the mean from bootstrap resampling. Points outside of plotting range are marked with Xs. *** *p* < 0.001, ** *p* < 0.01, * *p* < 0.05. FFA = fusiform face area, PPA = parahippocampal place area.

For decoding of face presence, classification accuracy was strong and widespread across occipitotemporal cortex for both the realistic and cartoon movies (Figure S5B). Classification accuracy in the ROIs (Figure 4B), was significant for the realistic movie in ventral occipital cortex (*M* = 0.678, CI = [0.653 to 0.701], *p* vs. 50% chance < 0.001), bilateral FFA (*M* = 0.628, CI = [0.593 to 0.661], *p* <0.001), and bilateral PPA (*M* = 0.635, CI = [0.601 to 0.666], *p* <0.001), as well as for the cartoon movie in ventral occipital cortex (*M* = 0.679, CI = [0.659 to 0.701], *p* < 0.001), bilateral FFA (*M* = 0.652, CI = [0.623 to 0.682], *p* <0.001), and bilateral PPA (*M* = 0.606, CI = [0.559 to 0.651], *p* <0.001). Again different from infants, cartoon movies did not yield significantly higher face decoding than realistic movies in ventral occipital cortex (*M* = −0.002, CI = [−0.025 to 0.022], *p* = 0.862), nor in bilateral FFA (*M* = 0.024, CI = [−0.010 to 0.058], *p* = 0.174) or bilateral PPA (*M* = −0.029, CI = [−0.061 to 0.002], *p* = 0.076).

For decoding of scene distance, classification accuracy was strong and widespread across occipitotemporal cortex for both the realistic and cartoon movies (Figure S5C). Classification accuracy in the ROIs (Figure 4C), was significant for the realistic movie in ventral occipital cortex (*M* = 0.509, CI = [0.481 to 0.542], *p* vs. 33% chance < 0.001), bilateral FFA (*M* = 0.422, CI = [0.400 to 0.446], *p* < 0.001), and bilateral PPA (*M* = 0.457, CI = [0.429 to 0.485], *p* < 0.001), as well as for the cartoon movie in ventral occipital cortex (*M* = 0.473, CI = [0.442 to 0.504], *p* < 0.001), bilateral FFA (*M* = 0.425, CI = [0.396 to 0.456], *p* < 0.001), and bilateral PPA (*M* = 0.410, CI = [0.371 to 0.451], *p* < 0.001). Whereas in ventral occipital cortex infants showed better decoding of scene distance for the cartoon movie, adults showed the opposite with significantly higher classification accuracy for the realistic movie (*M* = 0.036, CI = [0.004 to 0.069], *p* = 0.034). The same was true in bilateral PPA (*M* = 0.047, CI = [0.004 to 0.085], *p* = 0.038), but not bilateral FFA (*M* = 0.03, CI = [−0.032 to 0.028], *p* = 0.850).

## Discussion

Inspired by work mapping the optimal stimulus of visual cortex in adults and nonhuman primates (Hubel and Wiesel, 1962; Tanaka et al., 1991; Ponce et al., 2019), we assessed the preference of the infant visual system for exaggerated or ‘caricatured’ visual input by comparing fMRI responses to a matched pair of movies depicting the same narrative with photorealistic or cartoon visual styles. Cartoons led to greater synchrony and decoding in occipital cortex across infants. These findings largely replicated in separate awake infant fMRI datasets with realistic and cartoon movies that were not matched as carefully (Figure S6). Notably, the cartoon advantage was not found in young adults, suggesting that these results reflect a developmental process rather than some general stimulus difference between the movies.

An early preference for caricatured visual information may enhance the ability of infants to learn diagnostic features from their environment. Indeed, infant-directed speech helps infants map words onto their meanings (Ma et al., 2011) and infant-directed action promotes exploration (Meyer et al., 2023). Here, we found that more exaggerated input in the form of a cartoon enabled better decoding of faces and scenes, which are ecologically important visual features. Whether and how this enhanced decoding relates to better behavioral discrimination is not addressed by our study, but would be a clear prediction for future research. We did not find evidence of a cartoon advantage for feature decoding in adults; to the contrary, we found a double dissociation in ventral occipital cortex with better scene decoding for the realistic movie. Exaggerated input may thus provide a foundation for early learning that enhances later processing of more complex features. This idea is consistent with theories of perceptual development that emphasize the adaptive nature of limited early experiences with complex input (Turkewitz and Kenny, 1982; Vogelsang et al., 2024).

We have interpreted the greater reliability of neural responses in early visual cortex to the cartoon as reflective of perceptual tuning, reminiscent of stronger responses in primate visual cortex to the extremes of face space (Leopold et al., 2006; Freiwald et al., 2009; Koyano et al., 2021). However, neural synchrony can also relate to attentional engagement (Liu et al., 2025), such as heightened attention during the cartoon movie or greater consistency in the deployment of attention. We are unable to adjudicate this possibility in the present study, although we did not find effects in attentional regions, differences in BOLD signal from the eyeballs (Frey et al., 2021) (Figure S3), nor differences in the proportion of timepoints infants looked at the movie. This latter null effect, as well as in the proportion of timepoints retained after motion exclusion, help ensure that key comparisons between movies were not confounded by differences in data quality or quantity.

This study leaves open questions about the precise factors that drive neural preferences for more exaggerated visual content in the infant (but not adult) brain. In most infant-directed speech and action studies to date, various components (e.g., longer pauses, greater repetition, more exaggerated pitch/movements) are manipulated together, rather than isolated. We took a similar approach here, in the first study of its kind in the visual domain, with the cartoon manipulating several perceptual features at once. An exploratory analysis suggested that color and motion complexity alone could not explain enhanced neural synchrony in infants (Figure S2B), with equivocal results for shape complexity. Generative models may help isolate the role of caricaturing specific features by creating stimuli that are matched on some dimensions while differing on others (Hirschstein et al., 2025). They may also allow features to be manipulated more continuously, as when mapping the optimal stimulus of visual regions in nonhuman primates (Leopold et al., 2006; Freiwald et al., 2009; Koyano et al., 2021).

The movies used in this study depicted interactions between exotic animals in a simulated outdoor environment unlikely to have been experienced by our infant population first-hand. Thus, it is an open question whether the same cartoon advantage would emerge for familiar environments. One might expect weaker effects, if exaggerated content provides a scaffold for learning and is thus particularly beneficial for novel information. Alternatively, these effects may be insensitive to familiarity, as in language where infants show preferences for infant-directed speech in both native and non-native languages (Birulés et al., 2025). Importantly, we replicated our effects in a supplementary dataset of cartoon and realistic movies that depicted interactions among humans (Figure S6). Although these movies were not as carefully matched as those in the main dataset, the results suggest that our effects may reflect a more general, rather than stimulus-specific, property of the infant visual system.

We tested the sensory exaggeration hypothesis at two vastly different points in brain development (infants and adults) leaving the developmental trajectory of these effects unclear. The cartoon advantage may be limited to early life, a period of rapid visual changes (Braddick and Atkinson, 2011). Even among infants, we included a wide age range (4 to 15 months). This allowed us to consider the role of visual acuity, which may have still been developing in the youngest but not the older infants in the sample. If lower acuity was responsible for the sensory exaggeration (e.g., because of enhanced processing of high-contrast and coarser features), we would expect the difference between cartoon and realistic movies to be larger in the younger infants. Inconsistent with this possibility, there were no effects of infant age on neural synchrony or feature decoding in any of the regions that showed a cartoon advantage (Figure S7). At the other end of the spectrum, adults not only failed to show a cartoon advantage, but in fact showed the opposite for scene decoding in ventral occipital cortex and PPA, suggesting that extensive experience and perceptual learning may confer enhanced processing of realistic content.

In sum, we provide evidence consistent with the sensory exaggeration hypothesis — which proposes that developing organisms preferentially respond to exaggerated sensory input — in the visual system of human infants. It is tempting to speculate that children’s media, rife with exaggerated animated content (Wass and Smith, 2015), may be leveraging this neural preference. The parallels with other modalities (e.g., infant-directed speech and action) may point to a domain-general preference for caricatured input in infants, helping them build a scaffold for cognitive and perceptual development. How these early neural preferences for exaggerated input shape developmental outcomes may inform ways to best support attention and learning in early life.

## Acknowledgments

We are thankful to the families of infants who participated. We also acknowledge the hard work of the Yale Baby School team, including J. Daniels and K. Armstrong for recruitment, scheduling, and administration. Thank you to J. Fel, J. Cross, A. Püsök, M. Guha, and S. David, for help with gaze coding, R. Watts for technical support, and P. Deliwalla for sharing visual complexity code.

