## Supplementary Information for "Enhanced processing of cartoons in infant visual cortex"

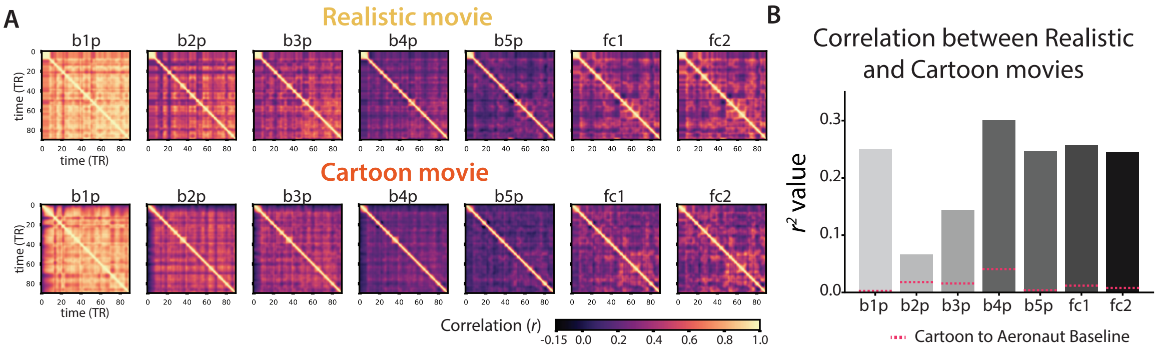


**Figure S1. Representations of realistic and cartoon movies in a convolutional neural network.** (A) Timepoint-by-timepoint correlation matrices for the pattern of unit activations in each layer of the VGG-19 network for the realistic (top) and cartoon (bottom) movies. (B) Representational similarity of the two movie styles for each layer of the network in terms of variance explained between the two corresponding correlation matrices in subpanel A after excluding the same and immediately adjacent timepoints (i.e., the diagonal and first 4 off-diagonals). The dashed red lines provide an across-movie null baseline obtained by comparing the cartoon version of The Lion King to “Aeronaut”, an unrelated cartoon of the same duration previously used in infant fMRI research (Yates et al., 2022).

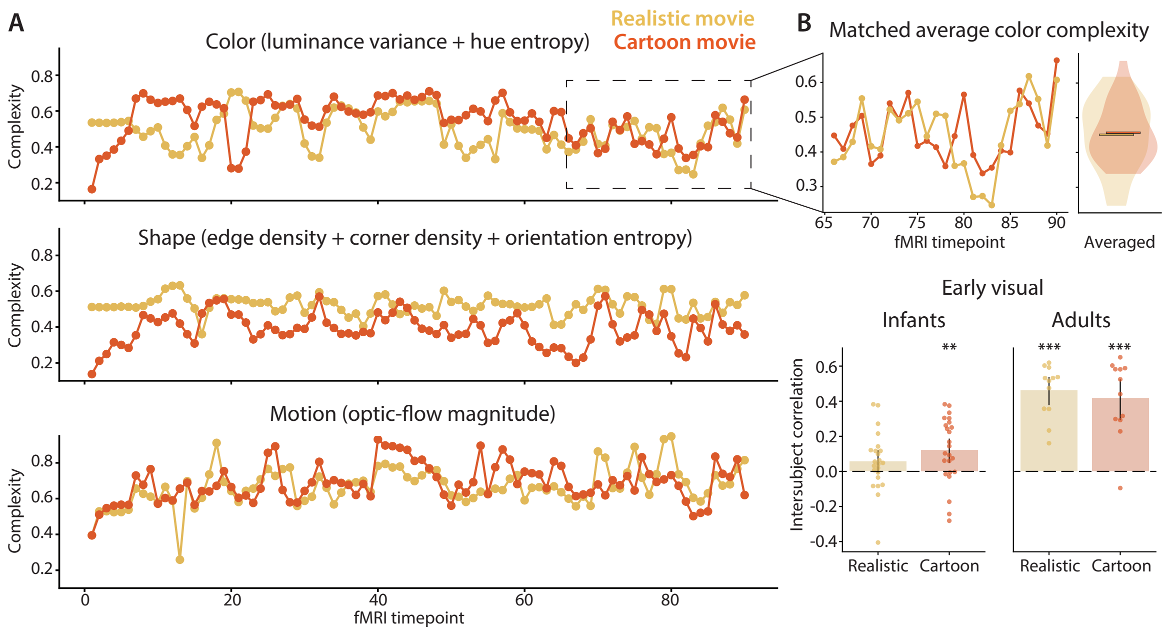


**Figure S2. Visual complexity in realistic and cartoon movies.** (A) Timecourse of different complexity measures for the realistic (yellow) and cartoon (orange) movies. (B) Although color complexity was generally higher for the cartoon versus the realistic movie, the values were approximately matched (and not statistically different) for the 25 timepoints (50 s) at the end of the movie (realistic: $M$ = 0.451, $SD$ = 0.100; cartoon: $M$ = 0.457, $SD$ = 0.081; difference: $M$ = 0.006, CI = [-0.045 to 0.058], bootstrap $p$ = 0.896). We recalculated intersubject correlation (ISC) in this subsegment for the early visual cortex ROI that showed enhanced neural synchrony for the cartoon vs. realistic movie in infants but not adults. We replicated the pattern of results from the entire movie, though effects were weaker likely because of the smaller sample of timepoints used to assess ISC. One infant could not be analyzed because only 2 timepoints were usable in the final subsegment of the realistic movie after motion and gaze exclusions. *** $p < 0.001$, ** $p < 0.01$.

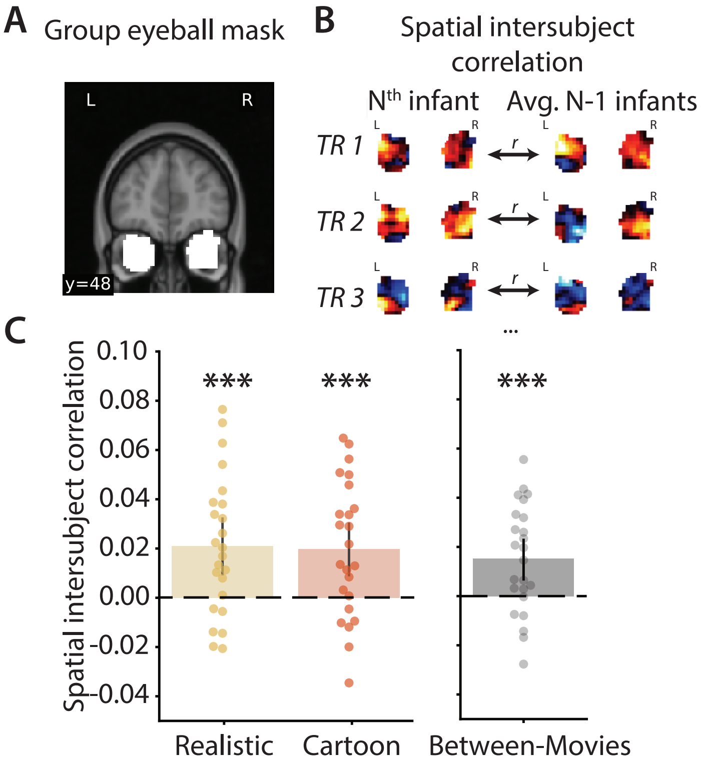


**Figure S3. Synchrony across infants in an eyeball ROI.** (A) Group eyeball ROI mask consisting of the common voxels across manually defined eyeball ROIs from each infant. (B) Spatial ISC was calculated by correlating patterns of voxel activity from bilateral eyeball ROIs for one held-out infant with the average of all other infants, separately for each TR and then averaged across TRs, repeated holding each infant out once. To maintain fine-grained patterns in the eyeballs and reduce signal influence from nearby frontal cortex, no spatial smoothing was performed during preprocessing for this analysis. (C) Spatial ISC values were significant in the eyeballs for both the realistic and cartoon movies in infants. There was no significant difference in spatial ISC between movies ($M$ = 0.001, CI = [-0.017 to 0.014], $p$ = 0.880). Moreover, there was similarity in the patterns of activity in the eyeballs between the two movies, calculated as the spatial ISC between an individual participant from one movie (e.g., cartoon) and the average of all other participants from the other movie (e.g., realistic; $M$ = 0.015, CI = [0.006 to 0.023], $p$ <0.001). Since patterns of activity in the eyeballs have been used for MR-based eye-tracking (Frey et al., 2021), this is consistent with equally reliable and similar gaze patterns across participants for the two movies. Dots represent individual participants and error bars represent 95% CIs of the mean from bootstrap resampling. *** $p$ < 0.001.

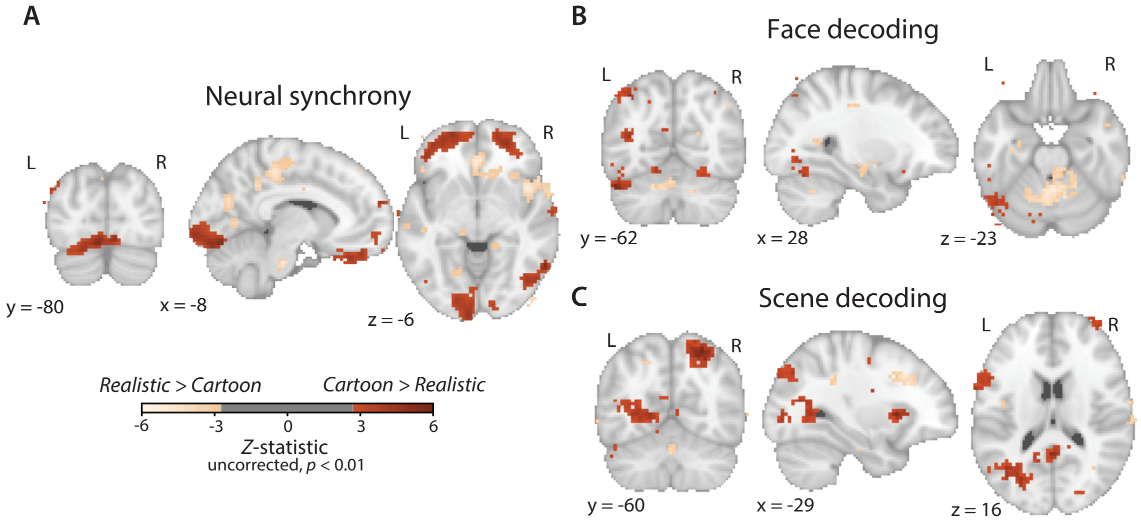


**Figure S4. Whole-brain comparison of neural synchrony and feature decoding between movies in infants.** Voxel-wise contrast of (A) intersubject correlation (ISC) values for the cartoon vs. realistic movie and (B, C) classification accuracy for the cartoon vs. realistic movie when decoding face presence and scene distance, respectively. Clusters are visualized at $p$ <0.01 (uncorrected), with peach indicating stronger values for the realistic movie and red indicating stronger values for the cartoon movie.

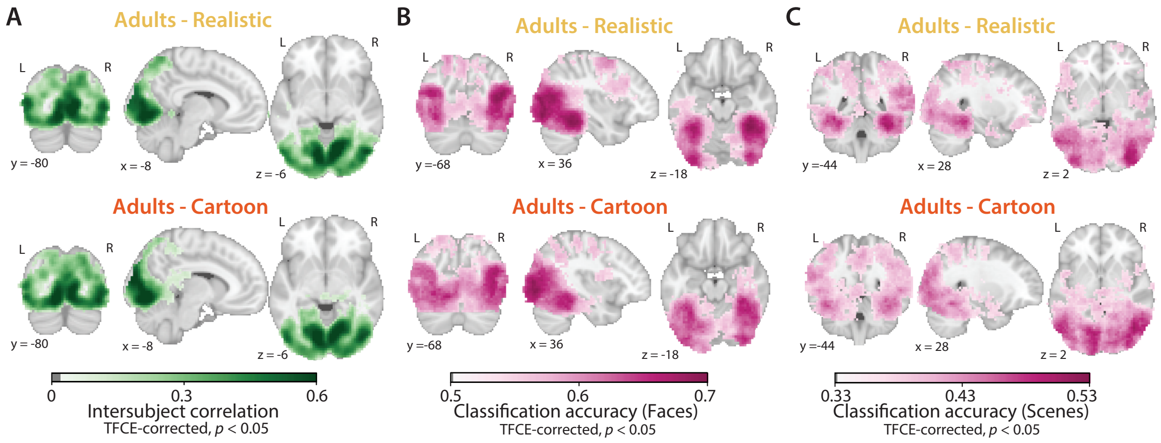


**Figure S5. Whole-brain neural synchrony and decoding for realistic and cartoon movies in adults.** (A) Voxel-wise intersubject correlation (ISC) values for the realistic and cartoon movies, thresholded at $p$ <.05, corrected with TFCE. Searchlight results for the decoding of (B) face presence and (C) scene distance. Voxels significant at $p$ <0.05 (TFCE corrected) are colored by the average classification accuracy across participants. Note that the color bar ranges differ slightly from Figures 2 and 3 to highlight the dynamic range of values in the adult data.

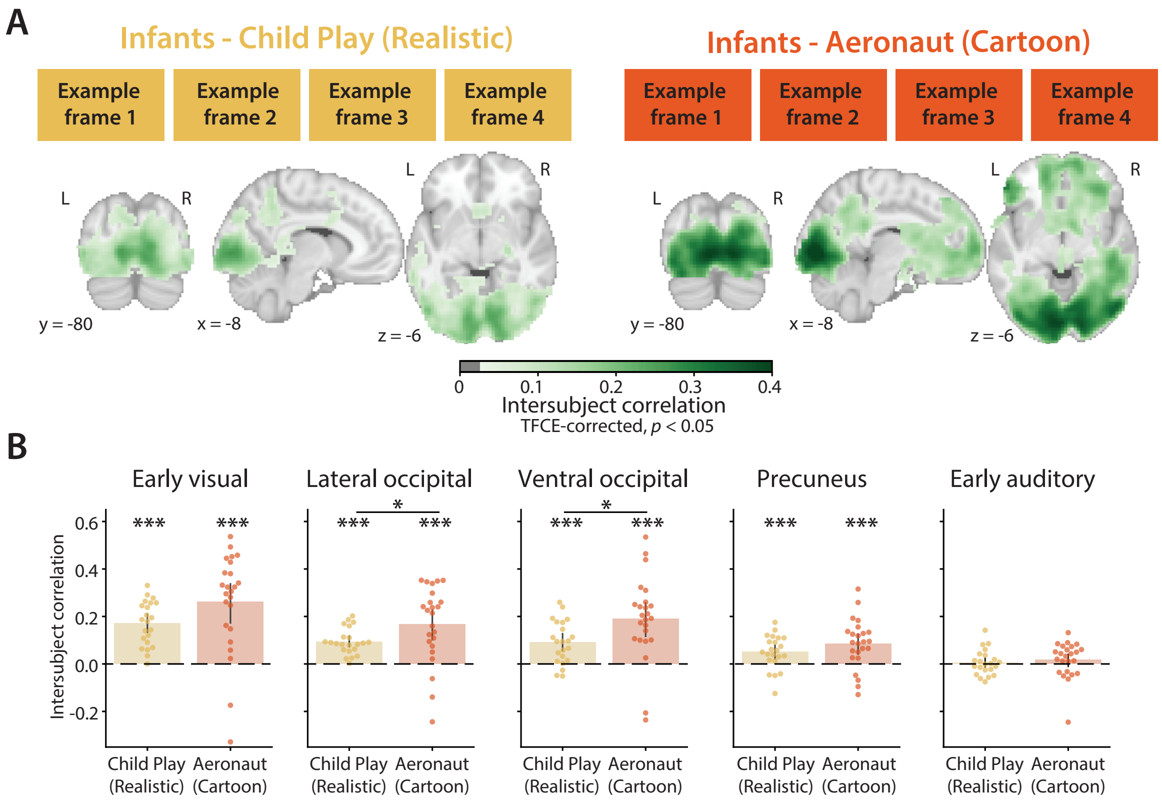


**Figure S6. Neural synchrony across infants in previously collected movie fMRI datasets not matched for content.** (A) Voxel-wise intersubject correlation (ISC) values for previously collected realistic (N = 22 infants, "Child Play"; (Ellis et al., 2025)) and cartoon movies (N = 24 infants, "Aeronaut"; (Yates et al., 2022)), thresholded at $p$ <.05, corrected with TFCE. Example movie frames containing faces are temporarily omitted to comply with bioRxiv policy. (B) ISC was significant in all three visual ROIs and the precuneus for both the realistic (early visual: $M$ = 0.171, CI = [0.134 to 0.211], $p$ <0.001; lateral occipital: $M$ = 0.094, CI = [0.071 to 0.117], $p$ <0.001; ventral occipital: $M$ = 0.091, CI = [0.054 to 0.128], $p$ <0.001; precuneus: $M$ = 0.051, CI = [0.016 to 0.078], $p$ <0.001; early auditory: $M$ = 0.006, CI = [-0.016 to 0.029], $p$ <0.600) and cartoon movies (early visual: $M$ = 0.261, CI = [0.173 to 0.340], $p$ <0.001; lateral occipital: $M$ = 0.168, CI = [0.103 to 0.228], $p$ <0.001; ventral occipital: $M$ = 0.190, CI = [0.120 to 0.259], $p$ <0.001; precuneus: $M$ = 0.086, CI = [0.047 to 0.128], $p$ <0.001; early auditory: $M$ = 0.018, CI = [-0.014 to 0.045], $p$ <0.230), with significantly higher values for the cartoon movie in lateral occipital ($M$ = 0.074, CI = [0.003 to 0.139], $p$ = 0.038) and ventral occipital ( $M$ = 0.099, CI = [0.019 to 0.176], $p$ = 0.026), and a trend in early visual ($M$ = 0.090, CI = [0.007 to 0.186], $p$ = 0.064); there was no significant difference between movies in the precuneus ($M$ = 0.035, CI = [-0.015 to 0.088], $p$ = 0.134) or early auditory ($M$ = 0.012, CI = [-0.030 to 0.048], $p$ = 0.510). Dots represent individual participants and error bars represent 95% CIs of the mean from bootstrap resampling. *** $p$ < 0.001, * $p$ < 0.05.

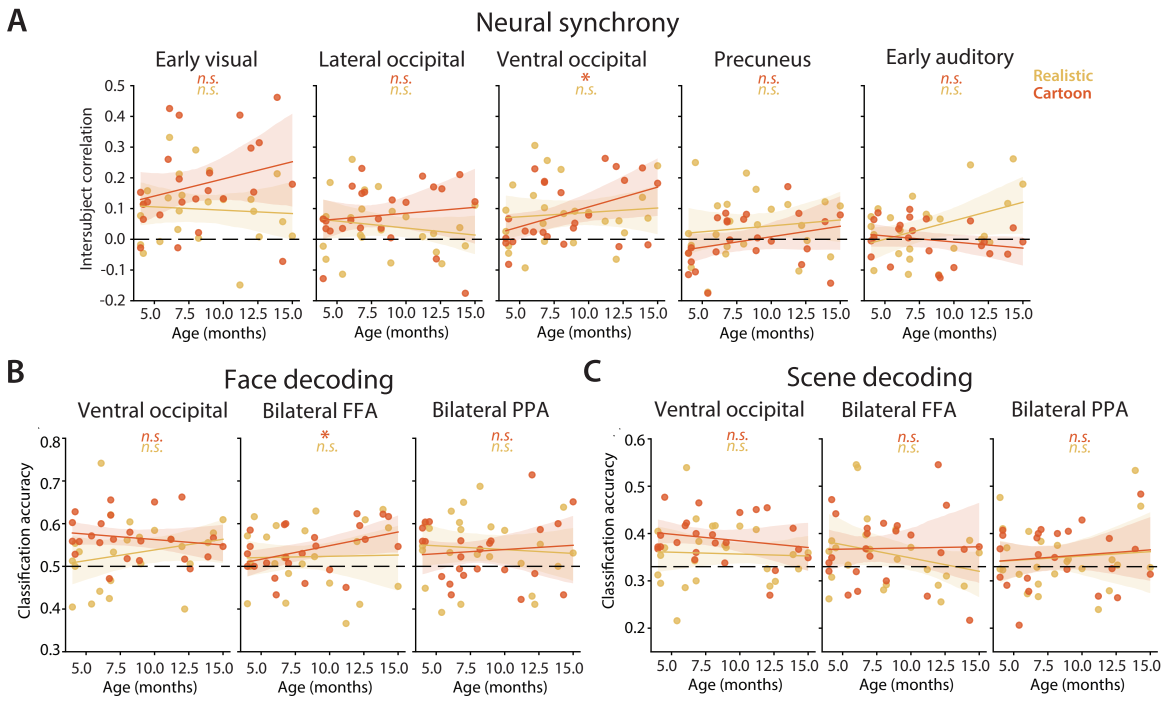


**Figure S7. Age effects within the infant sample.** (A) There was no evidence of a relationship between ISC and infant age in months for either the cartoon or realistic movie (*p*s >0.140), with the exception of a positive correlation for the cartoon movie in ventral occipital cortex ($r$ = 0.436, $p$ = 0.046) and a trend for the realistic movie in early auditory cortex ($r$ = 0.416, $p$ = 0.052). Dots represent individual participants and error band represents 95% CIs from bootstrap resampling. There was no evidence of a relationship between (B) face decoding or (C) scene decoding infant and age in months for either the cartoon or realistic movie (*p*s >0.120), with the exception of a positive correlation for face decoding of the cartoon movie in the FFA ($r$ = 0.403, $p$ = 0.038). * $p < 0.05$. FFA = fusiform face area, PPA = parahippocampal place area.
